# Armed to the teeth: the underestimated diversity in tooth shape in snakes and its relationship to feeding constraints

**DOI:** 10.1101/2022.12.13.520215

**Authors:** Marion Segall, Céline Houssin, Arnaud Delapré, Raphaël Cornette, Joshua Milgram, Ron Shahar, Anthony Herrel, Maïtena Dumont

## Abstract

Teeth are one of the most studied hard tissues in vertebrates. Their structure, composition and shape are related to dietary specialization in many species. At first glance, snake teeth all look similar; conical, sharp, curved. Yet, snakes, like other vertebrates, have very diverse diets that may have affected their shape. We compared the morphology of the teeth of 63 species that cover both the phylogenetic and dietary diversity of snakes. We predicted that prey properties play a role in shaping snakes teeth along with their feeding behavior. Limblessness combined to the peculiar feeding behavior of snakes impose strong functional constraints on their teeth, especially during arboreal or aquatic feeding. Our results show that prey hardness, foraging substrate and the main feeding constraint are drivers of tooth shape, size, and curvature. We highlight two main morphotypes: long, slender, curved with a thin layer of hard tissue for snakes that need a good grip on their prey and short, stout, less curved teeth in snakes that eat hard or long prey. Our study demonstrates the diversity of tooth morphology in snakes and the need to investigate the underlying functional implications to better understand the evolution of teeth in vertebrates.

## Introduction

Teeth allow most vertebrates to acquire food –and therefore energy—from their environment [1]. As such, they are one of the interfaces between an animal and its environment and their evolution depends both on intrinsic and extrinsic constraints. Teeth play different roles, from food acquisition to food processing, and have different functions (e.g., cutting, crushing, grinding, piercing). These functions depend on both food properties and/or food processing behavior and are related to tooth shape (e.g., [2]). Work by Crofts and colleagues (2020), based on pioneering studies by Lucas (2004)[1] and Massare (1987)[3], showed how tooth morphology is associated with prey properties and dental biomechanics (e.g., toughness, bending abilities, mode of failure). Rounded teeth, for example, allow to crush hard prey items, they are tough, can barely bend and are susceptible to fragmentation [2]. Because of their tight and reliable relationship with diet and their abundance in the fossil record, teeth have been suggested to be good indicators of past climate and paleoenvironments (e.g., [4]), and are used to make inferences on the ecology of extinct species [3,5–7]. By extension, tooth morphology could also be used to infer the feeding habits of secretive species that are sometimes only known from museum collection specimens; providing the link between tooth shape and food properties has been established. While most dental morphology studies have focused on mammals, because they benefit from a large variety of diets and teeth shape [8–10], a significant amount of work has been done on non-mammalian vertebrates [11]. Yet, quantitative comparisons of tooth morphology and its link to dietary ecology, in a phylogenetically and ecologically broad sample of species remain scarce in non-mammalian vertebrates. In this study, we investigated the relationship between dietary ecology and tooth shape in a group of non-model vertebrates: snakes.

Among vertebrates, macrostomatan snakes are peculiar as they are the only taxon able to ingest prey larger than their head without processing it. This behavior is related to an extraordinary organization of the skull that has become highly kinetic. Indeed, snakes must coordinate the movements of 8 pairs of cranial bones to catch, subdue, manipulate, and swallow their prey [12,13]. Despite the complexity of their feeding behavior, snakes have independently adopted a wide variety of dietary preferences (insects, mammals, birds, crustaceans) providing an opportunity to highlight convergences in their feeding apparatus [14]. In addition to constraints related to the physical properties of their food items, some feeding behavior may impose high loads on snake teeth, such as eating live and vivid preys, with or without the support of a solid substrate. These various mechanical challenges may have driven the evolution of tooth shape in snakes. Despite their richness and complexity in shape [15,16], studies on snake tooth morphology are scarce and either lack of a quantitative approach or are phylogenetically limited [11,17–20]. Fangs, and mostly front fangs, have recently attracted some scientific attention [21–26]. Yet, fangs are phylogenetically and functionally limited as their only purpose is to puncture the prey to deliver venom under the skin, and they are two highly derived teeth of over a hundred (for some species) that are involved in the whole feeding sequence.

Snake teeth are usually described as pointy and curved [11], they would therefore be considered as “piercing” specialists in the classification scheme as described in [2], and be related to a restricted diet composed of soft invertebrates and small fish. Yet, as previously mentioned, snakes show a broad variety of diets, but also a wide variety of feeding behaviors that involve their teeth such as ‘chewing’ [27], ripping [28–30], slicing [12,31], or swallowing without piercing (e.g. *Dasypeltis sp*.). Snake teeth are also involved in the whole feeding sequence, from prey capture to swallowing. Yet, the diversity of tooth morphology and function in snakes remains mostly under-explored. Here, we quantified and compared the dentary tooth morphology of 63 species that cover the phylogenetic and ecological breadth of snakes and tested four hypotheses relating their shape to feeding ecology:

- As for other vertebrates, tooth shape in snakes may be related to prey hardness, with hard-prey specialists having short and stouter teeth to limit failure [2].
- Because they swallow their prey whole and because they are vulnerable to predators during manipulation and swallowing, snakes must reduce the time and energy spend during feeding [32]. The overall shape of prey (bulky or elongated) involves different mechanical constraints [33]: bulky prey (e.g., anurans, mammals) require extensive manipulation [34] and repositioning, thus involving long, curved, and sharp teeth that are able to get a good grip on the prey. Elongated prey (e.g., snakes, eels), however, may require more ‘pterygoid walks’ (i.e., protractions and retractions of left and right tooth rows in alternating fashion assuring intraoral transport). In the latter case, short teeth that barely penetrate the prey may be more advantageous to avoid the cost of embedding and extracting the tooth rows.
- The mechanical constraints related to the feeding sequence in snakes may be dependent on the type of substrate in which it occurs. While feeding on the ground provides a solid substrate for the snake to support either itself or the prey during capture, subduction, manipulation and swallowing, aquatic and arboreal foraging do not. Aquatic predators face several constraints; first, their own buoyancy and that of their prey, second, the hydrodynamic forces that oppose any motion from the predator or the prey [35,36]. Arboreal manipulation and swallowing usually happen with the snake hanging from a branch, head hanging down, with no substrate support and almost without the help of the rest of its body. This behavior increases the chances of dropping the prey which may not be possible to catch again, unlike in an aquatic or terrestrial substrate. Thus, arboreal feeding requires both behavioral adjustments [37] and most probably long and sharp teeth that allow a good grip on the prey.
- These factors are not mutually exclusive (e.g., eel-eaters eat elongated prey under water) and can also be too restrictive for some species, such as terrestrial viperids that can either envenomate, release their prey and then track it, while other snakes hold on to the prey after striking which may impose high forces on their teeth [38]. We attributed to each species a main feeding constraint according to both the type of prey and its feeding behavior. This last factor may allow to highlight convergences between constraints, such as short and stout teeth for both hard and long prey or long and thin teeth for species that hold on to their prey, regardless of feeding substrate.

We dissected the dentary bone and used micro-CT scanner to obtain high resolution scans of the teeth which size varied between about 7mm to less than 1mm. We then used 3D geometric morphometrics to compare both the external and internal shape of the teeth. Shape information on inner part of the teeth allowed us to compare the thickness of the teeth in addition to shape. We also measured the curvature of each tooth. Then used phylogenetic comparative methods to test our predictive ecological factors to draw the link between teeth shape variation and dietary constraints in snakes.

## Material and Methods

### Tooth shape acquisition

We dissected 63 species that cover both the phylogenetic and dietary diversity of Alethinophidians (Supplementary Material 1 & 2). Our sample is composed of adult or subadult specimens from museum collections (AMNH, MNHN, Jerusalem University) and private collections (details in Supplementary Material 1 & 2). The dentary bone of the specimens was extracted, and CT scanned using the X-ray μCT-scanner Phoenix Nanotom S (General Electric, Fairfield, CT, USA) at the Institut de Genomique Fonctionelle, Ecole Normale Superieure (Lyon, France). The dentary with its teeth was scanned with a voltage of 100kV and a current of 70μA, for a voxel size between 0.97-7.50μm. The 3D reconstruction was done using Phoenix datos|x2 (v2.3.0, General Electric, Fairfield, CT, USA) and the subsequent segmentation was performed using VGStudioMax (v1.0, Volume Graphics GmbH, Heidelberg, Germany). We chose one tooth per specimen that was not broken and was not in the process of being replaced which was easily noticeable on the scans, through resorption of the bony attachment in most species. Our sample is composed of 63 dentary tooth (1 per species). We used 3D geometric morphometrics to get the 3D shape of the teeth. We placed 14 anatomical landmarks: 7 on the outer surface of the teeth, 7 on the pulp cavity surface (inner part of the tooth Supplementary Material 3) and 100 curve semi-landmarks (50 on each layer) using the software MorphoDig 1.2 [39]. Curves correspond to the anterior and posterior edge of both layers and to the limit of the tooth insertion into the bone. In addition, 42 and 65 surface semi-landmarks were respectively placed on the inner and outer surfaces of the teeth to obtain an accurate 3D representation of the tooth shape (Fig. 1). This template allows to obtain information both on their shape and on the thickness of the hard tissue material (dentine and enamel tissue). We placed the anatomical landmarks and curve semi-landmarks by hand on each specimen and checked for the repeatability of the positioning by digitizing these six times in five specimens for which the teeth looked similar. A Principal Component Analysis demonstrated the repeatability of the positioning of our 14 anatomical landmarks and the necessity of using curve and surface semi-landmarks (Supplementary Material 4). We used the ‘Morpho’ package [40] to project and relax the surface semi-landmarks on each specimen and to slide the curve and surface semi-landmarks while minimizing the bending energy between the specimen and the template [41]. We then performed a Procrustes superimposition using the function *gpagen* of the ‘geomorph’ package [42], the resulting Procrustes coordinates were used to test our hypotheses. The projection, relaxation and Procrustes superimposition were performed using R version 3.4.4 [43].

**Fig. 1:**
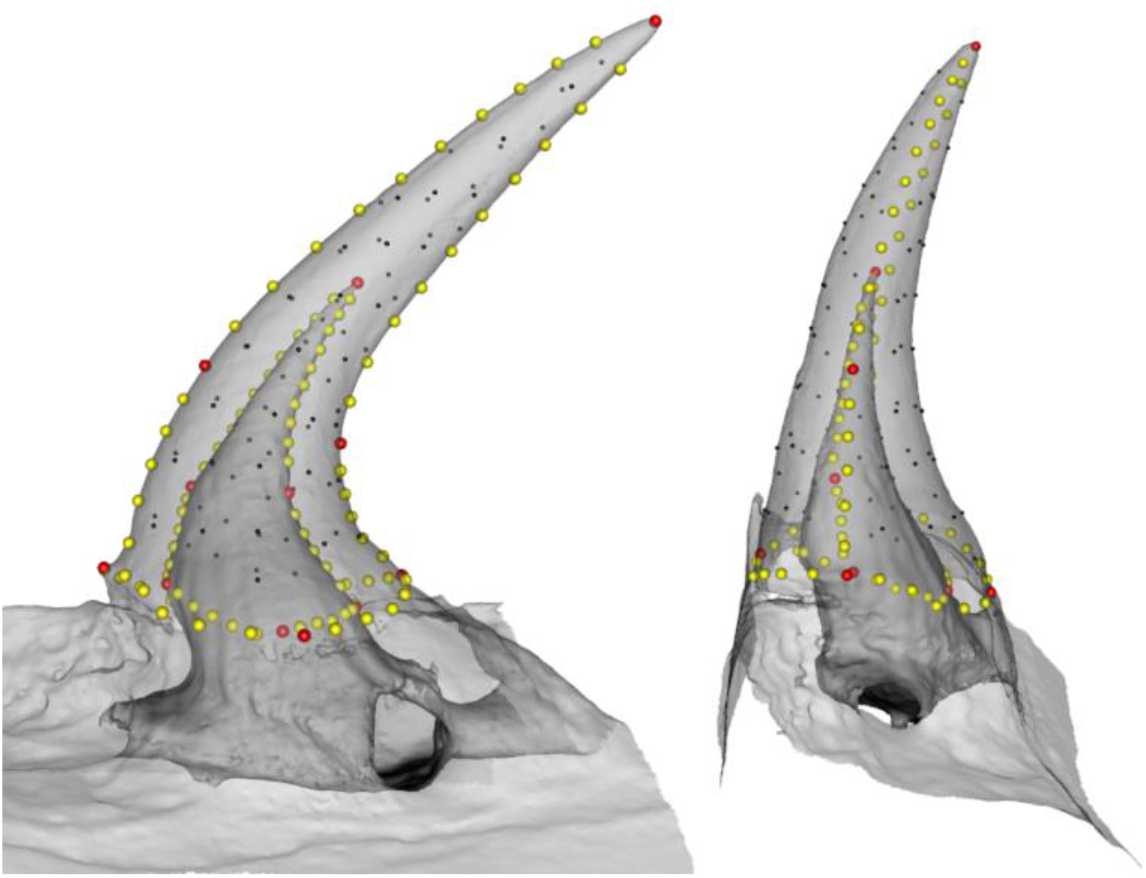
Snake tooth template illustrating the landmarks used for geometric morphometric analysis. **3D mesh of a dentary** tooth of *Natrix tessellata (HUJI 16537)* in medial (left) and anterior (right) view. The scan is semi-transparent to show the inner layer of the tooth. Landmarks are represented by spheres colored in red for anatomical landmarks, yellow for curve semi-landmarks and black for surface semi-landmarks. A total of 221 landmarks were used to describe the 3D shape of snake teeth.

As 3D geometric morphometrics remove size information, we took linear and angle measurements of the teeth to test their potential link with dietary ecology. We measured the length of the curvature (LC) along with maximal and mean curvature (κ_mean_, κ_max_) of each teeth using the FIJI (v1.53q) plugin ‘Kappa’ [44]. We used a snapshot of the medial side of each tooth, obtained in GeomagicStudio 2013 (3D Systems, Rock Hill). We digitized the midline to obtain its global curvature (see examples in Supplementary Material 5, raw measurements available in Supplementary Material 1). From the mean and max curvature measurements, we calculated the corresponding degree of curvature (D_Cmean_, D_Cmax_) using the following equation:

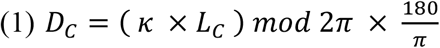

### Dietary constraints

An extensive bibliographic study was performed to characterize the dietary constraints related to each of the 63 species considered in this study (Supplementary Material 1). Following our hypotheses, we defined four factors that could impact the shape of the teeth in snakes: prey hardness, prey shape, feeding substrate and main dietary constraint. These factors are related to both the main prey item and the feeding behavior which we thoroughly reviewed for the 63 species (Supplementary Material 1). We defined the three levels of prey hardness: soft (e.g., gastropods, annelids, birds, mammals), medium (e.g., amphibians fish, thin-scaled lizards such as anoles) and hard (insects, snakes, hard-scaled lizards such as skinks, [45]). Prey shape has 2 levels: bulky or long following descriptions in [46]. Mammals prey were considered bulky as the hind limbs are particularly difficult to swallow and require extensive manipulation in snakes. For generalist snakes that do not show a preferred item or only have little information on their diet available, we did not attribute them to any prey shape group (“na” in Supplementary Material 1). The foraging substrate has 3 levels: ground, water and branch depending on where the swallowing of the food occurs. The main constraint encountered by snakes while feeding has five levels: hard (e.g., chitinous preys, hard-scaled lizards), long (e.g., snakes, soft-scaled lizards, earthworms), bulky (e.g., mammals, amphibians), hold (e.g., snakes that must maintain vivid preys), slippery (e.g., fish, snails, amphibian eggs). When several prey items were recorded for a species, we used the most commonly ingested prey.

### Analyses

We used phylogenetic comparative methods, using a tree pruned from Pyron & Burbrink (2014)[47], if species were not present in this tree, we used the closest relative (Supplementary material 2). We tested whether teeth curvature length, mean degree of curvature and maximum degree of curvature were correlated with each other using a Pearson correlation test (function *cor.test* from the ‘stats’ package). We then tested whether these measurements were associated with our factors using the function *phylANOVA* of the package ‘phytools’ using 1000 simulations and a Holm correction for the post hoc pairwise t-tests. The normality of the data distribution was checked using a Shapiro test and variables were log-transformed to ensure normality when necessary. We used the function *procD.pgls* in ‘geomorph’ [48] to run phylogenetic ANCOVAs to test link between tooth shape and our different dietary constraints, using the log-transformed centroid size of the tooth as a covariate. Since our predictive factors are not entirely independent from one another, we performed one model per factor. We compared the fit of our different models using the function *model.comparison* in the ‘RRPP’ package [49], that calculated the log-likelihood of each model. Statistical significance was tested by performing 10000 permutations. Subsequent post-hoc pairwise comparisons were done using the *pairwise* function in ‘RRPP’. All geometric morphometric, statistical analyses and visualizations were performed in R version 3.4.4 [43] (R code and data available in Supplementary Material), except the landmark acquisition performed in MorphoDig [39]. To highlight shape differences between ecological groups, we used mesh deformation of the template specimen toward the mean landmark configuration for each group, using the *plotRefToTarget* function of the ‘geomorph’ package. We also used the function *spheres3d* from the ‘rgl’ package that allows to visualize both the inner and outer surfaces of the teeth, providing information on teeth thickness (Fig. 5).

## Results

### Variation in tooth length and curvature

Dentary tooth length (LC) in our sample varies between 0.37 – 6.68mm, with a median of around 1.2mm (Fig. 2). Tooth length is related to the main feeding constraint (F=7.59, P=0.001), with snakes that must hold their prey, or that feed on bulky or slippery prey having longer teeth than snakes feeding on hard prey (P_hard/bulky_=0.003, P_hard/hold_=0.03, P_hard/slippery_=0.004). Prey hardness is also related to tooth length (F=11.67, P=0.0005), with snakes feeding on hard prey having shorter teeth than snakes preying upon softer prey (P_hard/medium_<0.001, P_hard/soft_=0.002; Fig. 2).

**Fig. 2:**
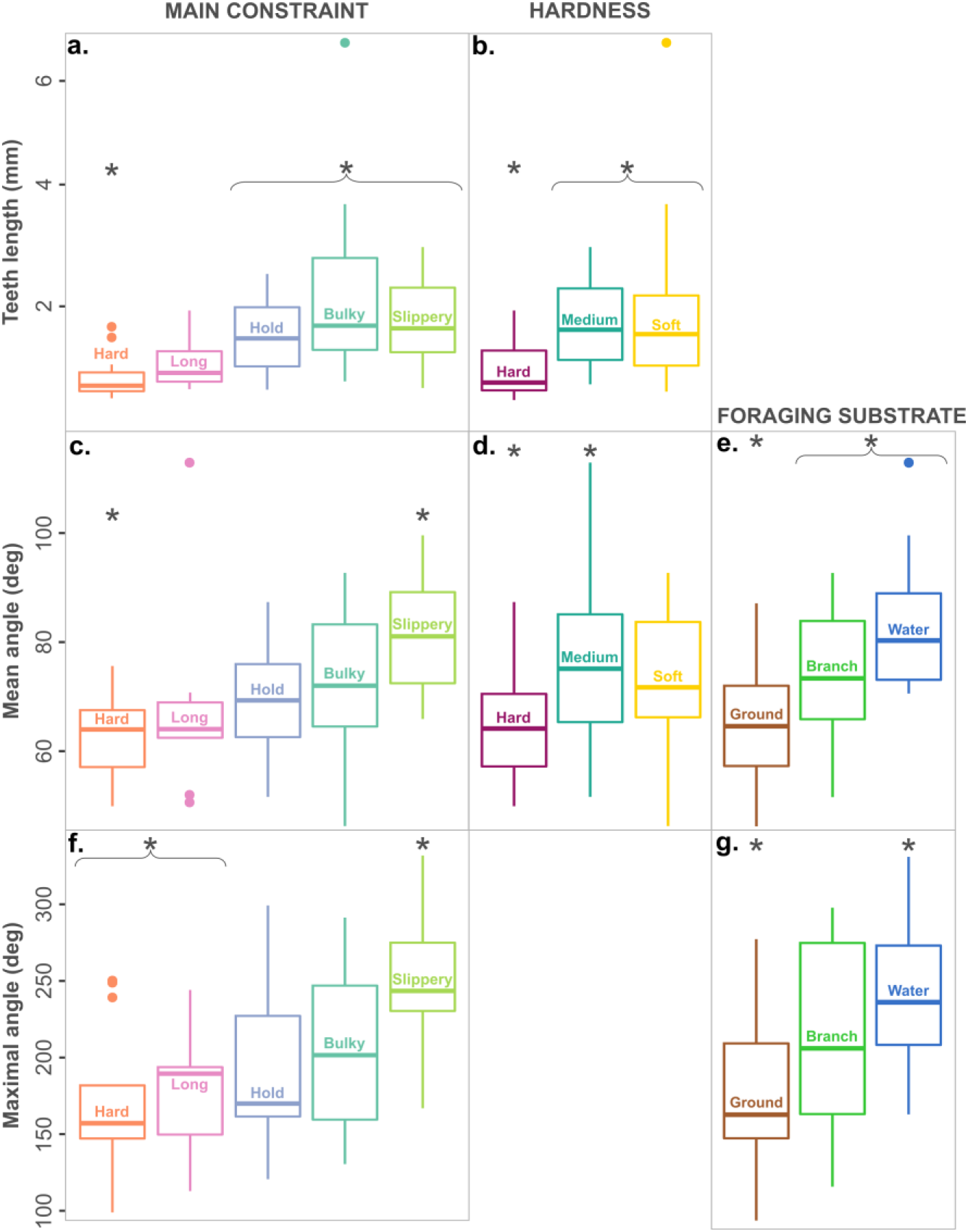
Boxplots representing the differences in tooth length (a, b), mean angle (c, d, e) and max angle (f, g) depending on feeding constraints. a-c-f: differences related to the main feeding constraint; b-d: the hardness of the prey and e-g: the foraging substrate. Statistically significant differences are indicated by *. Groups in the same brace are not significantly different but are different from groups with the asterisk (e.g., teeth length is different between hard versus hold, slippery and bulky).

The average curvature (mean angle) of the teeth varies between 46 – 113°, with a median of around 70.6°. It significantly differs depending on the main feeding constraints (F=4.03, P=0.036), prey hardness (F=4.8, P=0.04) and the foraging substrate (F=10.14, P=0.003) of species. Snakes feeding on slippery prey (P_slippery/hard_=0.01) and snakes feeding on prey with a medium softness (P_hard/medium_=0.01) have more curved teeth (larger mean angle) than snakes feeding on hard prey. Teeth of snakes that forage under water or on branches are significantly more curved than snakes feeding on the ground (P_ground/branch_=0.04, P_ground/water_=0.004; Fig. 2).

The maximal degree of curvature varies between 99 – 332° with a median around 194° and is significantly associated with the main feeding constraint (F=4.84, P=0.016) and the foraging substrate (F=6.62, P=0.013). Snakes feeding on slippery prey have a higher maximal angle of curvature than species feeding on hard (P_slippery/hard_=0.006) and long prey (P_slippery/long_=0.019; Fig. 2). Snakes feeding under water have significantly a higher maximal curvature than snakes feeding on the ground (P_ground/water_=0.017; Fig. 2).

Neither the length nor the degrees of curvature are significantly associated with the shape of the prey (all P>0.1). We found significant, yet weak, positive correlations between teeth length and the mean (P=0.03, R=0.26) and max degrees of curvature (P=0.03, R=0.27), but Figure 3 shows that snakes feeding on slippery and bulky prey tend to have longer and more curved teeth while snakes feeding on hard and long prey have shorter and straighter teeth. The correlation between the mean and max degrees of curvature is also significant; snakes with a higher mean curvature also have a high maximal angle of curvature (P<0.0001, R=0.64).

**Fig. 3:**
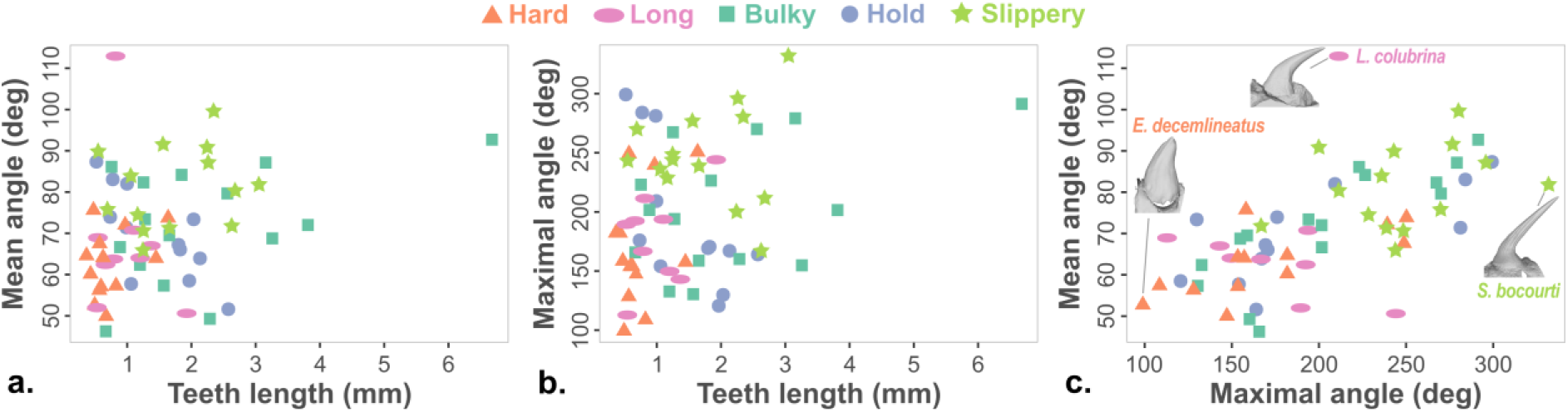
Correlations between linear measurements of the tooth. a. tooth length and mean angle, b. tooth length and maximal angle, c. mean angle and maximal angle. Each dot represents one species and is colored and shaped according to the main constraint encountered during feeding (legend above the graph).

### Tooth shape and feeding constraints

There is a large variation in the shape of the teeth: from very thin, slender and with a pulp cavity that runs along the full length of the teeth to short, stout teeth with a thick layer of tissue. This variation is represented along PC1 (Fig. 4). PC2 differentiates between curved teeth, with a large base and a short pulp cavity and stout, less curved teeth. In the morphological space, species that eat long or hard prey are mostly positioned on the left side of the plot while species that eat bulky, slippery prey and species that must hold their prey are mostly positioned on the right side, suggesting a relationship between tooth shape and the main feeding constraint. Our statistical analyses validate this (F_4,62_=2.17, P=7.10^−4^, R^2^=0.1) and pairwise post hoc tests show significant differences in tooth shape between snakes eating hard versus slippery prey and species that must hold their prey (P_hard/hold_=0.0012, P_hard/slippery_=0.0017), and between species eating long prey versus those that hold their prey and those that eat slippery prey (P_long/hold_=0.013, P_long/slippery_=0.004). Tooth shape is also associated with prey hardness (F_2,62_=3.38, P=2.10^−4^, R^2^=0.08), with differences between snakes feeding on hard prey versus medium and soft ones (P_hard/soft_=0.002, P_hard/medium_=0.0001). The foraging substrate also significantly impacts tooth shape (F_2,62_=3.29, P=3.10^−4^, R^2^=0.07), with species feeding on the ground being different from both arboreal and aquatic species (P_ground/tree_=0.0023, P_ground/water_=0.0025). Yet, as for tooth length and curvature, tooth shape is not significantly associated with prey shape. The model that best fits our data is the main constraint (AIC=-3880.836), then the foraging substrate (AIC=-3235.691), and prey hardness (AIC=-3217.872).

**Fig. 4:**
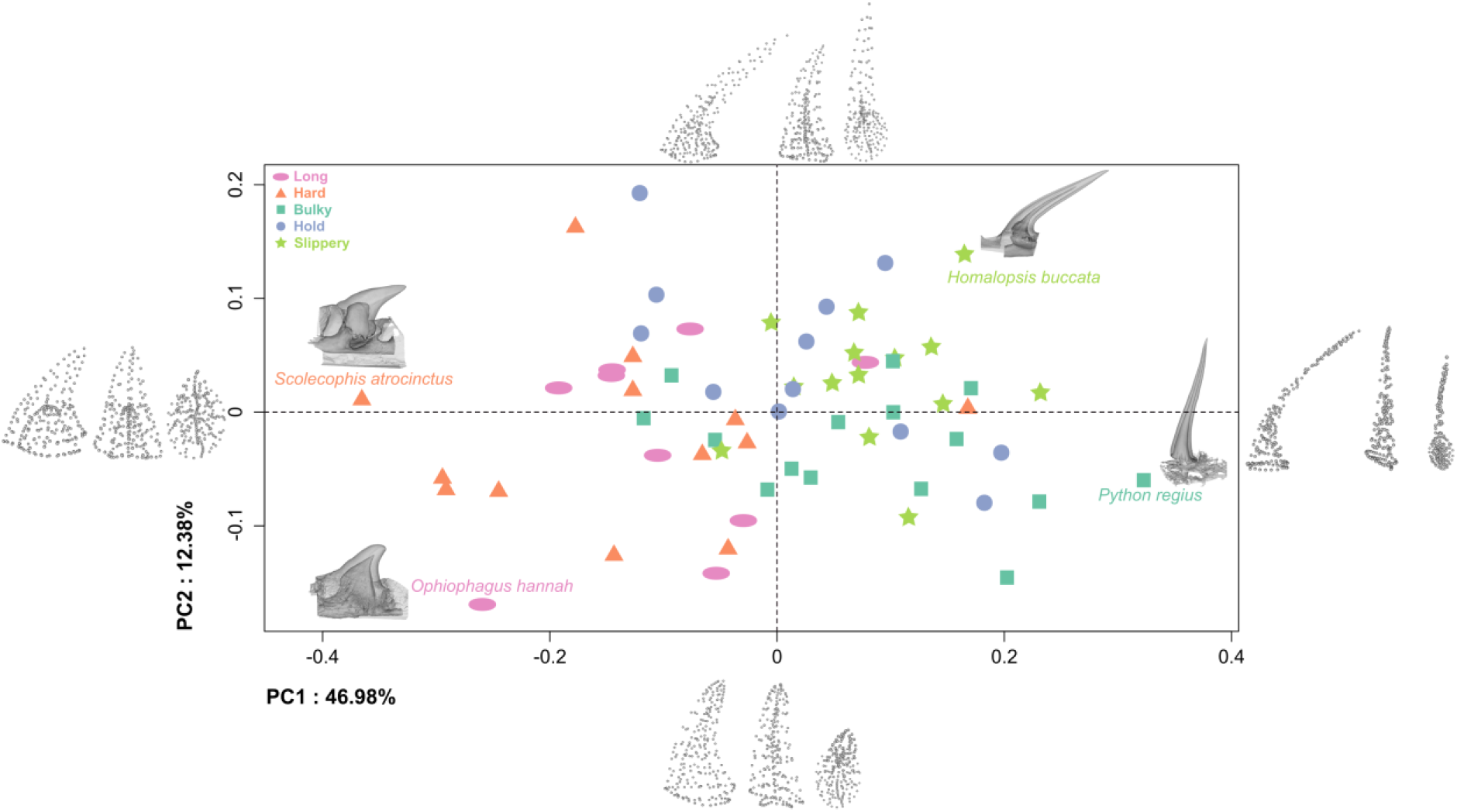
Tooth shape variability in snakes. Plot of the two first principal components that represent almost 60% of the shape variation. On the extreme of each PC axis is represented the associated landmark configuration in (from left to right) medial, anterior, and top view. Four examples of scans are positioned in the morphological space to highlight shape diversity. Each dot represents one species and is colored and shaped according to their main feeding constraint.

Overall, regardless of the predictive factor, the morphological variation between groups that differ significantly distinguishes between short, bulky, and thick teeth versus long, slender, and thin layered teeth (i.e., teeth with a pulp cavity running to the tip of the tooth) (Fig. 5). Snakes with slender teeth belong to the hold and slippery groups, they eat medium to soft prey without the help of a solid substrate (i.e., aquatic or arboreal feeding). Snakes that have robust teeth eat long or hard prey, typically on the ground (Fig. 5).

**Fig. 5:**
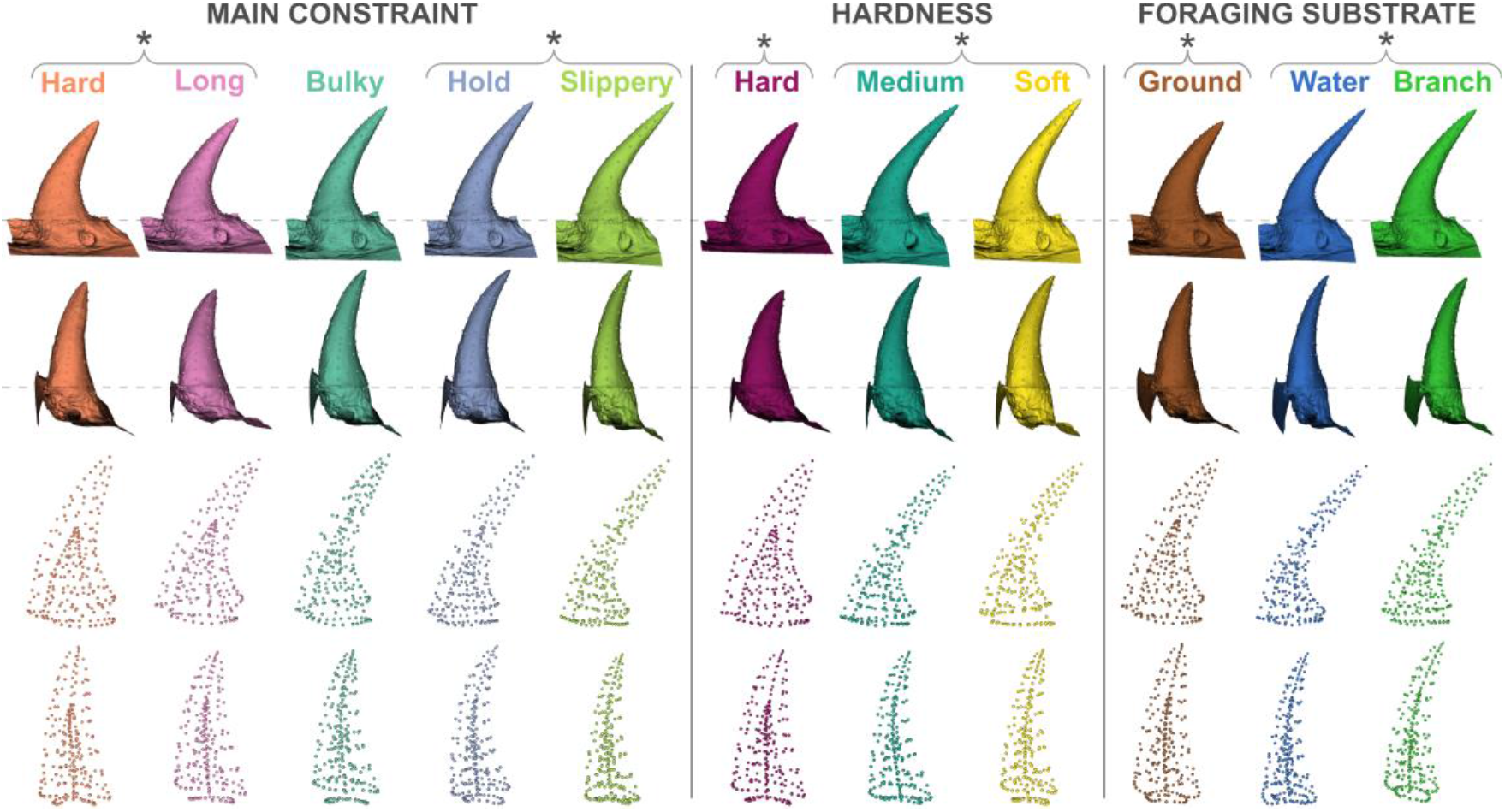
Mean teeth shapes associated with significant ecological predictors. First line shows the medial side of the teeth and second line the anterior view. Meshes were aligned according to the curve that delimits the base of the teeth (see dashed lines). The third and fourth lines show landmark configurations using spheres. Statistically significant differences are indicated using braces: groups that in the brace are not significantly different but are different from groups in the other brace (e.g., long, and hard are different from hold and slippery).

## Discussion

From the sharp, elongated and nearly straight teeth of the ball python (*Python regius*, Fig. 4), to the round, bulky and nearly flat teeth of the king cobra (*Ophiophagus hannah*; Fig. 4), the morphological variability of snake dentary teeth is far larger than what transpires from the literature. Yet, snake teeth remain easily identifiable among vertebrate as they share some characteristics; they are all conical, to some extent, with various degrees of sharpness and they are curved in two directions, posteriorly and medially, although to various degrees as well (Fig. 2–5). This curvature may prevent breakage of the teeth when the snake strikes at its prey at high velocity and high acceleration [19]. An increase in the curvature is associated with a decrease in the maximal stress undergone by the teeth when out-of-plane forces are involved, which is likely to be the case during a feeding sequence. The curvature may also allow the prey to slide over the teeth as far as possible into the mouth to be impaled when moving backwards, thus preventing a potential escape. The mean and maximal curvature are significantly related to tooth length, but these relationships are very weak and probably only significant because of the largest species in our sample (Fig. 3). However, the correlation between the average and maximal degree of curvature is interesting; it was not obvious that this relationship existed as the overall curvature can either be caused by an elbow-like shape (e.g., *Homalopsis buccata*, Fig. 4) or a continuous curvature (e.g., *Laticauda colubrina*, Fig. 3). In the first case, the maximum of curvature is high, but the mean curvature is low, while the reverse is true for the second case. Yet, our results show that more curved teeth also have a larger maximal angle of curvature and overall straighter teeth have a smaller maximal angle of curvature. These various degrees of curvature are associated with specific feeding constraints.

### Prey hardness

As predicted, snakes eating hard prey have short, stout and less curved teeth contrary to softer-prey eaters that have long, slender, and more curved teeth. During penetration of the prey, short and stout snakes teeth undergo a relatively smaller maximal stress compared to slender teeth, coming from the compression force and, the stress is concentrated at the tip of the tooth [20,50] which may prevent failure of the teeth while feeding on hard items but may cause fragmentation and wear of the tip of the tooth. We noticed some tooth fragmentation in species such as *Fordonia leucobalia*. *Fordonia* is a peculiar species that feeds on crustaceans and dismembers them with its mouth [51]. This behavior likely imposes high load on the teeth, thus aggravating the fragmentation. The dentine and enamel organization in this species is also unique in our sample and will be the described in another study. Apart from this species, most hard prey specialists do not use their teeth to dismember their prey but swallow them whole. Snake teeth seem disadvantageous to feed on hard prey, but unlike other durophagous vertebrates, snakes generally do not use their teeth to crush. They use their teeth to manipulate and swallow their prey, thus, they may not need to resist high loads. In fact, snakes feeding on hard prey generally do not embed their teeth in the prey. Some species even have independently evolved hinged teeth that fold back when they swallow the prey but may rise up if the prey move backwards preventing escape [52,53]. Some legless lizards specialized on eating snakes or hard scaled lizards show a similar specialization suggesting a convergent evolution of hard-prey specialists in squamates [54].

Snakes eating softer prey have longer and more slender teeth that they can embed in their prey to get a good grip during manipulation and to prevent the prey from escaping or falling from their mouth. The differences in tooth shape, size and curvature related to prey hardness do not distinguish between very soft prey, such as snails, earthworms, mammals, and less soft prey such as fish or small-scaled lizards. The major differences lie in the distinction between hard and softer prey, and this result can suggest two hypotheses. First, the adaptation of the teeth to related to hardness may not be a continuum following the spectrum of hard-to-soft. There may be a hardness threshold beyond which the teeth breaks but below which the teeth resist. Second, we probably need to measure prey hardness to be able to draw precise a relationship between food properties and teeth shape. As suggested in Crofts and colleagues (2020), food material properties such as stiffness (resistance to deformation) and toughness (energy absorption before failure) are important characteristics (they are both related to hardness) to fully understand a correlation between prey and tooth morphology. They are, however, difficult to measure and should be measured for each prey. Prey hardness clearly has a role in teeth evolution in snakes, but the lack of significant difference between softer prey categories suggests that other feeding constraints may also contribute to the diversity of teeth in snakes.

### Foraging substrate

We hypothesized that snakes feeding on the ground benefit from the support of a solid substrate to capture, manipulate and swallow their prey, unlike aquatic and arboreal specialists that need a tight grip on their prey. This hypothesis is supported by our results as the shape and the curvature of teeth are different between snakes that eat on the ground and the two other categories. Arboreal and aquatic feeders have relatively thinner and more curved teeth, while the terrestrial group has bulkier and less curved teeth (Fig. 2 & 5). Tooth length is not statistically different between ecologies and, despite not being significant, there are differences between the shape of arboreal and aquatic snakes that are worth noticing. Teeth of aquatically feeding snakes are more slender and curved, with a larger maximal degree of curvature, providing them with an “elbow-like” shape compared to arboreal snakes whose teeth are curved all along (Fig. 4 *Homalopsis buccata*, Fig. 5: *Laticauda colubrina, Subsessor bocourti*). The pulp cavity runs farther up in aquatic than in arboreal species. These two specificities may allow the teeth of aquatic snakes to be more flexible than arboreal snakes. This hypothesis remains to be tested but the teeth of aquatically feeding snakes are weakly ankylosed [45], suggesting another adaptation to prevent breakage of these slender and sharp teeth [54].

### Prey shape

Unlike other vertebrates that either reduce the size of their food by chewing, crushing, or cutting into smaller, easily ingestible pieces (e.g., most mammals) or swallow relatively small items (e.g., some birds, lizards), snakes can swallow large prey and are gape-limited predators. In snakes, prey shape is related to manipulation effort, and head and skull shape [33,46,55–58] but it does not significantly affect the size, curvature, or shape of their teeth. The shape of the prey may not be sufficient by itself to drive tooth morphology and some other factors may be more important. For instance, snakes that eat long prey sometimes specialize on eating long and hard prey (e.g., centipedes), long and soft (e.g., earthworms) or long prey in an aquatic context (e.g., eels) and the reverse is true for bulky prey; they can be soft (e.g., mammals) or hard (e.g., eggs). Thus, it is possible that, while the shape of prey plays a role in shaping other parts of the feeding apparatus, it is not constraining enough by itself to significantly impact tooth shape. An alternative explanation would be the presence of ‘true’ generalist, species for which we were not able to determine a dominant prey and that were considered as a proper group, which may bias our analyses.

### Main constraint

Among our four predictive factors, the main mechanical constraint related to the preferred prey item or the feeding behavior is the only one significantly linked to all our tooth characteristics (size, mean and max curvature, shape). Among the five categories, two are always significantly different: species that eat hard prey and species that eat slippery prey. The hard prey group is characterized by teeth that are short, stout, with the smallest mean and maximal curvature (Fig. 2 & 5). Their pulp cavity is the short and the relative thickness of hard tissue is larger than for the other groups, making their teeth more robust. On the opposite, slippery prey eaters are characterized by long, slender, and highly curved teeth (Fig. 2 & 5). Their pulp cavity is long and provides the teeth with a relatively thin layer of hard tissue which may allow more bending, but this hypothesis remains to be tested. Snakes that eat long prey tend to resemble the hard prey group (Fig. 5) and occupy the same area in the morphospace (Fig. 4). The tooth shape of snakes that must hold their prey is similar to that of slippery prey eaters (Fig. 5).

All our results highlight two morphotypes: long, thin, highly curved teeth with a thinner layer of hard tissue (long pulp cavity running along the teeth) versus short, stout, straighter with thicker layer of hard tissue (short pulp cavity). Long, slender teeth are associated with feeding constraints that require a good grip on the prey, whether the prey is slippery, or soft, or if there is no solid substrate to support and facilitate the limbless feeding sequence. Thin teeth undergo high stress during penetration into the prey, due to bending, and are more susceptible to failure [50]. Thin teeth are also highly affected by even the slightest axial force that cause high stress [20,50]. Thus, a certain amount of bending might be beneficial to avoid breakage, especially in a feeding context where the prey cannot be correctly restrained. Short and stout teeth are associated with hard or long prey which both impose either high and/or repeated loading on the teeth but the stout shape allows to decrease the maximal stress that originates from the compression force and concentrates it in the tip [20,50]. Rajabizadeh and colleagues (2020)[20] compared the mechanical properties associated with tooth shape in two sister species; one that eats hard prey and has short, stout teeth versus a generalist with a slenderer teeth. They used finite element analyses to compare von Mises stress and deformation during loading from various angles on the two teeth. They demonstrated that, as suggested by Bar-On’s results (2019)[50], snake teeth barely undergo any stress when applying a tangential force to the tip. Deviation of the applied force from the tangent of the tip imposes a higher and more widely distributed stress in the slender tooth than in the stout one. These results suggest that some of the morphological variation we highlight here seem to be related to mechanical adaptation of the teeth to dietary constraints. Yet, the two compared shapes are far from representing the large variability in teeth morphology in snakes and many functional aspects of snake teeth remain unexplored such as the effect of curvature or the effect of variation in the inner shape of the teeth on its biomechanical properties. Conclusions based on the teeth of other vertebrates are hardly applicable to snakes. Snake teeth fulfill different functions than those of other vertebrates; they play a major role in prey capture and intra-oral transport, but they are rarely used to reduce the size of prey items. Although snakes have acquired constraining diets such as durophagy, the function of their teeth is not to crush (except for *Fordonia*) but to transport the whole prey into the digestive tract. Therefore, the trade-off highlighted for the teeth of durophagous vertebrates, between convex teeth that reduce the force needed to break a hard item but increases the strain in the tooth versus concave tooth that reduces the strain but require higher forces to break the prey, does not apply to durophagous snakes [59]. Our study shows that prey mechanical properties are not the only drivers of tooth morphology, but feeding behavior, and more globally feeding ecology, impose a variety of constraints that impact their size and shape. Future investigations of the biomechanics of snake teeth may help draw the link between their morphological and behavioral variability and would enrich our understanding of tooth evolution and function in vertebrates. Experimental designs [21], simulations [20,50] and analytical tools [60] have recently been developed and can be used to better understand the dental biomechanics of snakes using the shapes highlighted in the present study, in a functionally relevant context.

## Conclusion

The relationship between dietary ecology and dental morphology has been quite well established in many vertebrate taxa [11]. Snakes are hyper-specialized which may have constrained the evolution of their tooth morphology and function. Here, we demonstrate that this is not the case; the morphology of snake teeth is diverse, and this diversity is associated with several aspects of their feeding ecology. Two main teeth shapes were highlighted in our study: short and robust and long and slender teeth. Long teeth are present in snakes that need a good grip on their prey, such as soft-bodied prey or feeding without a solid substrate. Short teeth are associated with hard and/or long prey item that usually do not involve penetration of the prey. This is the first study quantifying and comparing the morphology of snake teeth in a large sample of species with different ecologies. We hope this will open the way to further investigations on the underlying mechanical properties of snake teeth which would allow to better understand the adaptive nature of their morphological diversity. Finally, we only explored only one (the dentary bone) of the four tooth-bearing bones. Previous work has demonstrated that these bones are organized in functional modules [14]. It is possible that teeth morphology differs at the intra-individual level, depending on the role of the teeth in the feeding sequence. To date, there is only one study comparing the shape of teeth between bones in one generalist species [19]. Results show that there is some degree of intra-individual variation in this species, but a comparative study is needed to test whether this pattern apply to species with a more specialized diet.

## Acknowledgements

We thank the Gans Fund for funding this study. MS was also funded by the European Union H2020 – Marie Skłodowska-Curie Global Fellowship (GA101024700). Special thanks to the people and institutions who provided specimens to enrich our sample and without whom this study would not have been possible: Anthony Herrel (MNHN), David Kizirian and Lauren Vonnahme from the herpetological collection of the American Museum of Natural History (New York), Nicolas Vidal from the herpetological collection of the Museum National d’Histoire Naturelle (Paris), Marc Herbin (MNHN), Boaz Schacham from the herpetological collection of Jerusalem, Rémi Ksas and Antoine Planelles (Venomworld), Ya-Wei Li, Vincent Prémel, Ludovic Faure, Yoan Eynac and Karine Falco. We thank Mathilde Bouchet from ENS Lyon for her time helping to set the specimens and teaching MS to use the CT scanner, and Adrien Izzet for his explanations about curvature and angles.

## Availability of data and material

The Rdata and Rcode will be provided in the Supplementary Material and the 3D scans will be uploaded in MorphoSource.

## Authors’ contributions

MS and MD conceived and led the research project. MS, CH, AD help collecting the data. MS led the writing of the manuscript. RC, RS, JM, AH helped in writing the grant proposal. All authors contributed to the interpretation and discussion of the results, and to the editing of the manuscript.

