## Supplementary Material for "Armed to the teeth: the underestimated diversity in tooth shape in snakes and its relationship to feeding constraints"

**Supplementary Material 1:** List of specimens used in this study along with information about the collection, scans, feeding ecology and teeth measurements. Teeth length= $L_C$ (curvature length), mean angle = $D_{Cmean}$ (average degree of curvature), max angle = $D_{Cmax}$ (maximal degree of curvature).

| **Species** | **Collection** | **Scan resolution (µm)** | **Diet** | **Main constraint** | **Shape** | **Hardness** | **Foraging substrate** | **Tooth length (mm)** | **Mean angle (º)** | **Max angle (º)** | **References** [1] |
|---|---|---|---|---|---|---|---|---|---|---|---|
| *Anilius scytale* | A. Herrel | 1.87 | reptiles, amphibians | long | long | hard | ground | 1.926 | 50.6 | 244.1 | [2,3] |
| *Calabaria reinhardtii* | A. Herrel | 1.42 | mammals | bulky | bulky | soft | ground | 1.278 | 73.4 | 194.0 | [4] |
| *Candoia carinata* | A. Herrel | 2.62 | reptiles | hold | long | hard | branch | 1.800 | 67.2 | 169.3 | [5] |
| *Eryx jaculus* | HUJI 3634 | 2.00 | mammals | bulky | bulky | soft | ground | 1.847 | 84.2 | 226.5 | [6,7] |
| *Boa constrictor* | Yoan Eynac | 7.50 | endotherm generalist | bulky | bulky | soft | branch | 6.678 | 92.7 | 291.4 | [8] |
| *Corallus annulatus* | A. Herrel | 2.75 | endotherm generalist | hold | bulky | soft | branch | 1.824 | 66.1 | 170.6 | [8,9] |
| *Cylindrophis ruffus* | Ludovic Faure | 1.75 | reptiles, amphibians | long | long | hard | ground | 1.369 | 67.0 | 143.0 | [10–15] |
| *Python regius* | A. Herrel | 2.15 | endotherm generalist | bulky | bulky | soft | ground | 2.288 | 49.3 | 160.2 | [16] |
| *Morelia spilota* | A. Herrel | 3.37 | mammals | bulky | bulky | soft | ground | 3.813 | 72.0 | 201.5 | [17] |
| *Acrochordus javanicus* | A. Herrel | 2.27 | fish | slippery | long | medium | water | 2.628 | 71.8 | 166.9 | [18] |
| *Acrochordus granulatus* | A. Herrel | 1.59 | fish | slippery | long | medium | water | 1.659 | 71.3 | 238.7 | [18–21] |
| *Xenodermus javanicus* | A. Herrel | 1.00 | fish | slippery | long | medium | water | 0.695 | 75.8 | 269.8 | [22] |
| *Aplopeltura boa* | Karine Falco | 1.47 | snails | slippery | long | soft | branch | 1.248 | 65.9 | 243.9 | [23–25] |

| | | | | | | | | | | | |
|---|---|---|---|---|---|---|---|---|---|---|---|
| *Pareas carinatus* | Anthony Herrel | 1.22 | snails | slippery | long | soft | branch | 0.551 | 89.8 | 243.1 | [26–29] |
| *Eristicophis macmahoni* | Latoxan | 2.00 | ectotherm generalist | hard | long | hard | ground | 1.638 | 73.7 | 250.2 | [30,31] |
| *Daboia russelii* | Venomworld | 2.12 | generalist | bulky | na | medium | ground | 2.559 | 79.7 | 270.0 | [32] |
| *Causus sp.* | Ludovic Faure | 1.59 | amphibians | bulky | bulky | medium | ground | 0.757 | 86.1 | 223.0 | [33,34] |
| *Echis leucogaster* | Latoxan | 1.75 | arthropods, mammals | hard | long | hard | ground | 0.593 | 57.1 | 153.5 | [35,36] |
| *Bitis gabonica* | Latoxan | 3.77 | mammals | bulky | bulky | soft | ground | 3.156 | 87.1 | 279.1 | [37] |
| *Tropidolaemus wagleri* | AMNH R50991 | 1.64 | mammals | hold | bulky | soft | branch | 1.966 | 58.5 | 120.6 | [38] |
| *Gloydius halys* | Ludovic Faure | 1.18 | mammals | bulky | bulky | soft | ground | 0.889 | 66.7 | 201.6 | [39] |
| *Bothriechis schlegelii* | Venomworld | 2.57 | generalist | hold | na | medium | branch | 2.575 | 51.6 | 164.0 | [1,40] |
| *Agkistrodon piscivorus* | Venomworld | 2.00 | generalist | bulky | na | medium | ground | 1.656 | 69.5 | 158.8 | [41–45] |
| *Crotalus sp.* | A. Herrel | 4.25 | mammals | bulky | bulky | soft | ground | 3.259 | 68.8 | 154.8 | [46] |
| *Subsessor bocourti* | A. Herrel | 2.75 | fish | slippery | long | medium | water | 3.051 | 81.8 | 331.8 | [10,47] |
| *Cantoria violacea* | LKC ZRC 2.3317 | 1.00 | crustaceans | hard | long | hard | ground | 0.969 | 72.0 | 239.2 | [47,48] |
| *Fordonia leucobalia* | LKC ZRC 2.3294 | 1.59 | arthropods | hard | bulky | hard | ground | 0.827 | 57.3 | 108.5 | [48–50] |
| *Gerarda prevostiana* | LKC ZRC 2.3292 | 1.00 | arthropods | hard | bulky | hard | ground | 0.625 | 64.1 | 153.4 | [47–49] |
| *Homalopsis buccata* | A. Herrel | 1.70 | fish | slippery | long | medium | water | 2.263 | 87.1 | 295.9 | [10,47,51] |

| | | | | | | | | | | | |
|---|---|---|---|---|---|---|---|---|---|---|---|
| *Malpolon insignitus* | HUJI 16560 | 1.81 | generalist | bulky | na | medium | ground | 1.570 | 57.3 | 130.5 | [52–54] |
| *Atractaspis engaddensis* | HUJI 16567 | 1.75 | reptiles, amphibians | hard | long | hard | ground | 0.430 | 60.1 | 181.8 | [55] |
| *Micrurus psyches* | A. Herrel | 0.97 | snakes | long | long | hard | ground | 0.545 | 68.9 | 112.8 | [56] |
| *Ophiophagus hannah* | Latoxan | 2.27 | snakes | long | long | hard | ground | 1.203 | 64.0 | 149.7 | [57,58] |
| *Dendroaspis viridis* | Latoxan | 1.50 | mammals | hold | bulky | soft | branch | 0.987 | 71.4 | 281.2 | [59] |
| *Naja annulata* | Latoxan | 2.50 | fish | slippery | long | medium | water | 2.682 | 80.3 | 211.4 | [60] |
| *Naja nigricollis* | Latoxan | 1.50 | generalist | bulky | na | medium | ground | 1.199 | 62.4 | 132.7 | [61] |
| *Laticauda colubrina* | AMNH R38111 | 1.06 | fish | long | long | medium | water | 0.819 | 112.9 | 77.3 | [50,62–67] |
| *Aipysurus laevis* | Latoxan | 2.37 | fish | slippery | long | medium | water | 2.244 | 90.8 | 200.0 | [50,64,65,68, 69] |
| *Hydrophis platurus* | A. Herrel | 1.92 | fish | slippery | long | medium | water | 2.347 | 99.6 | 280.2 | [50,68,70–77] |
| *Grayia ornata* | A. Herrel | 1.29 | fish | slippery | long | medium | branch | 1.559 | 91.5 | 276.7 | [60,78] |
| *Boiga dendrophila* | Latoxan | 1.83 | generalist | hold | na | medium | branch | 2.134 | 63.9 | 167.1 | [79] |
| *Boiga cynodon* | Latoxan | 2.00 | birds | hold | bulky | soft | branch | 2.034 | 73.4 | 129.9 | [79] |
| *Dasypeltis scabra* | A. Herrel | 1.04 | eggs | hard | bulky | hard | ground | 0.572 | 67.5 | 248.8 | [80] |
| *Dispholidus typus* | Latoxan | 1.75 | birds, lizards | hold | bulky | medium | branch | 1.062 | 57.7 | 154.1 | [81] |
| *Philothamnus semivariegatus* | Latoxan | 2.00 | lizards | hold | bulky | hard | branch | 0.736 | 74.0 | 176.0 | [59,78,82] |

| | | | | | | | | | | | |
|---|---|---|---|---|---|---|---|---|---|---|---|
| *Eirenis decemlineatus* | HUJI 4780 | 1.50 | arthropods | hard | bulky | hard | ground | 0.495 | 52.6 | 98.9 | [83,84] |
| *Eirenis lineomaculatus* | HUJI 16485 | 1.75 | arthropods | hard | bulky | hard | ground | 0.475 | 75.6 | 158.1 | [83,84] |
| *Oxybelis aeneus* | A. Herrel | 1.12 | generalist | hold | na | medium | branch | 0.998 | 82.0 | 209.2 | [85,86] |
| *Scolecophis atrocinctus* | A. Herrel | 1.01 | arthropods | hard | bulky | hard | ground | 0.368 | 64.6 | 181.8 | [87] |
| *Coluber constrictor* | AMNH R27367 | 1.25 | insect, snakes, lizard | hard | long | hard | ground | 1.449 | 64.0 | 157.1 | [88–90] |
| *Stenorrhina degenhardtii* | AMNH R38087 | 1.59 | arthropods | hard | bulky | hard | ground | 0.567 | 56.2 | 128.0 | [91–93] |
| *Gonyosoma boulengeri* | A. Herrel | 1.50 | endotherm generalist | hold | bulky | soft | branch | 0.776 | 83.1 | 284.0 | [1] |
| *Coronella girondica* | A. Herrel | 1.40 | lizards | long | long | hard | ground | 0.786 | 63.8 | 166.7 | [94] |
| *Lampropeltis triangulum* | AMNH R177123 | 1.22 | mammals | bulky | bulky | soft | ground | 0.668 | 46.3 | 165.7 | [95] |
| *Natrix tessellata* | HUJI 16537 | 1.50 | fish | slippery | long | medium | water | 1.165 | 74.5 | 228.5 | [96–98] |
| *Liodytes rigida* | AMNH R177031 | 1.00 | arthropods | hard | bulky | hard | ground | 0.672 | 49.9 | 147.2 | [99–102] |
| *Heterodon nasicus* | Yoan Eynac | 1.30 | reptiles, toads | bulky | bulky | hard | ground | 1.255 | 82.3 | 267.4 | [103–106] |
| *Imantodes cenchoa* | Vincent Prémel | 1.00 | reptiles, amphibians | hold | long | hard | branch | 0.521 | 87.3 | 299.3 | [3,107] |
| *Atractus flammigerus* | MNHN 80-24 | 1.25 | earthworms | long | long | soft | ground | 1.107 | 70.8 | 193.7 | [3,108,109] |
| *Sibon sp.* | A. Herrel | 1.02 | amphibian eggs, earthworms, snails | slippery | long | soft | branch | 1.058 | 83.9 | 236.1 | [110,111] |

| | | | | | | | | | | | |
|---|---|---|---|---|---|---|---|---|---|---|---|
| *Helicops sp.* | A. Herrel | 1.20 | fish | slippery | long | medium | water | 1.252 | 70.6 | 248.5 | [3,112–114] |
| *Siphlophis compressus* | Vincent Prémel | 1.59 | lizards | long | long | medium | ground | 0.663 | 62.5 | 192.4 | [1,3] |
| *Clelia clelia* | MNHN 79-47 | 1.40 | reptiles | long | long | hard | ground | 0.529 | 52.0 | 189.5 | [1] |

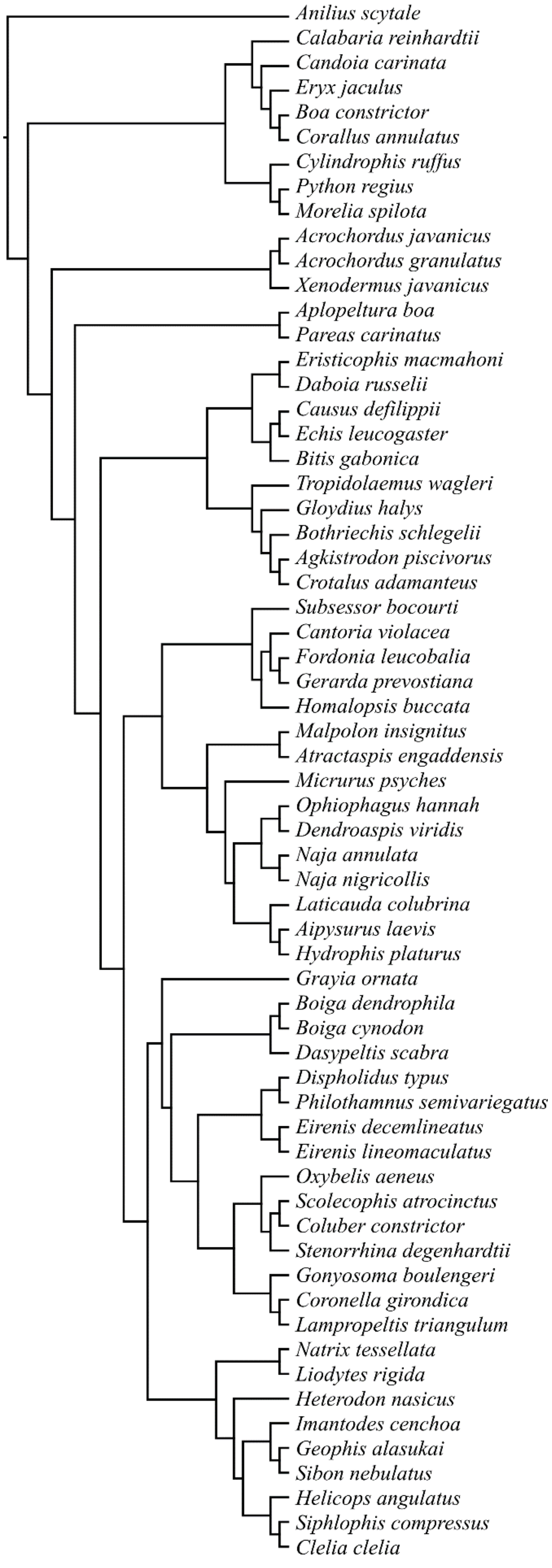

**Supplementary Material 3:** Description of anatomical placed on the inner (blue) and outer (red) layers of the teeth. Left figure: medial view of the tooth. Right picture: posterior view of the tooth.

| LM# | Name |
|---|---|
| 1 | Tip of the pulp cavity |
| 2 | Most anterior point of the tooth insertion |
| 3 | Most posterior point of the tooth insertion - according to the orientation with the vascular canal |
| 4 | Most medial point of the tooth insertion |
| 5 | Most lateral point of the tooth insertion |
| 6 | Max. curvature - anterior |
| 7 | Max. curvature - posterior |
| 8 | Tip of the tooth |
| 9 | Most anterior point of the tooth insertion |
| 10 | Most posterior point of the tooth insertion - according to the orientation with the vascular canal |
| 11 | Most medial point of the tooth insertion |
| 12 | Most lateral point of the tooth insertion |
| 13 | Max. curvature - anterior |
| 14 | Max. curvature - posterior |

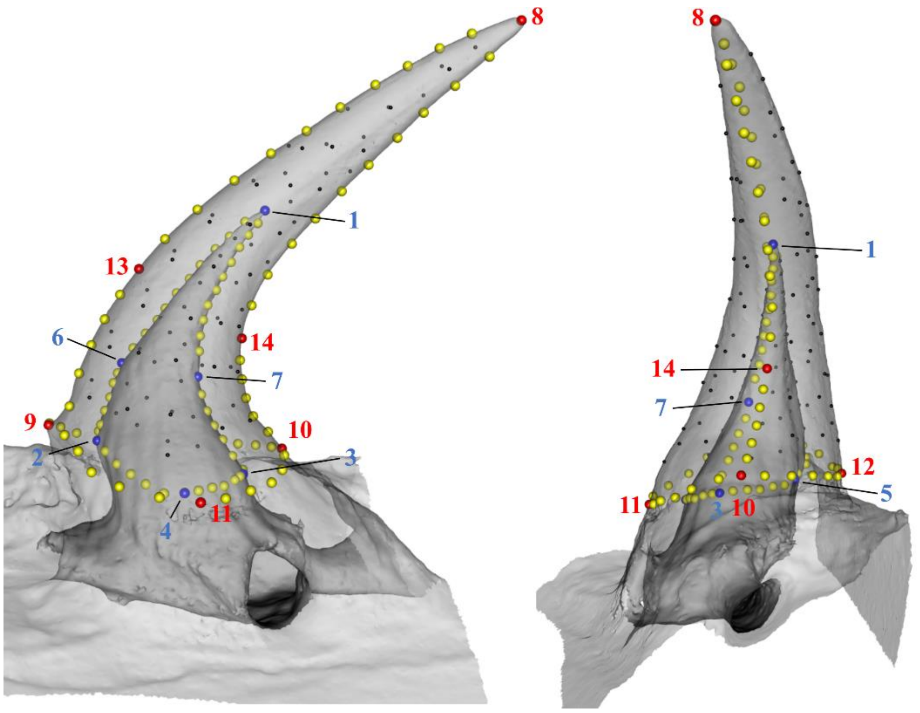

**Supplementary Material 4:** PCA plot showing the repeatability of the anatomical landmark positioning. We landmarked teeth that appear very similar. The plot shows that, despite being similar in shape, repetitions of the same specimen group together and are separated from the other specimens, based on the 14 anatomical landmarks. Yet, some specimens are close (*Candoia carinata* "C_car" and *Boiga cynodon* "B_cyn"), thus suggesting a need for more accurate shape information using curve and surface semi-landmarks.

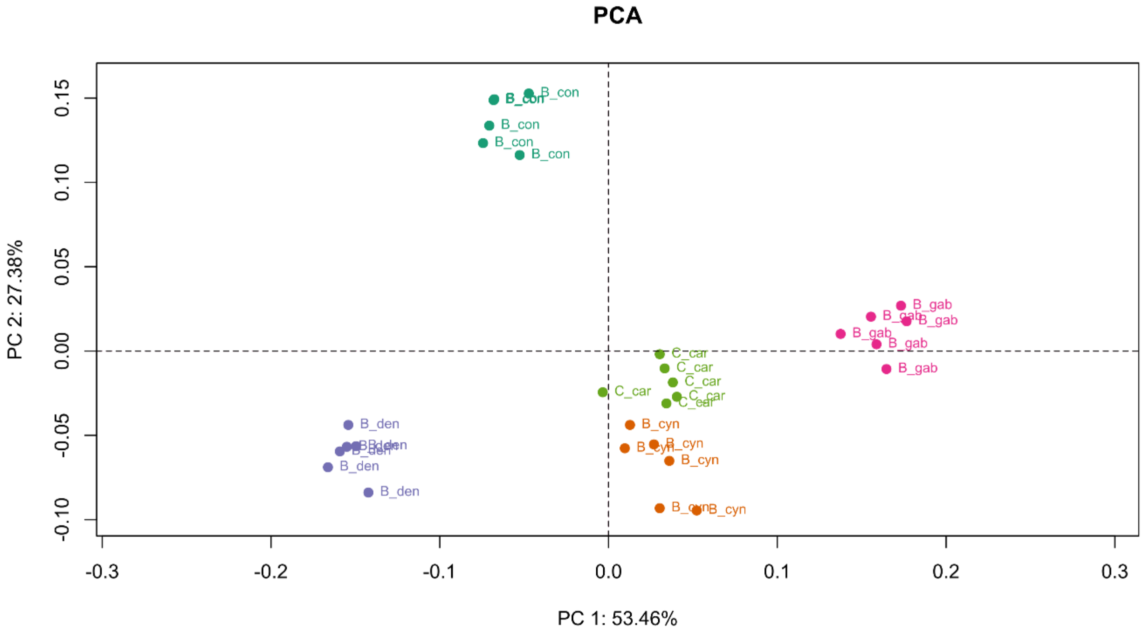

**Supplementary Material 5:** Two examples of curvature digitization using the plugin Kappa in FIJI: *Boa constrictor* (top snapshot), *Eirenis decemlineatus* (bottom snapshot).

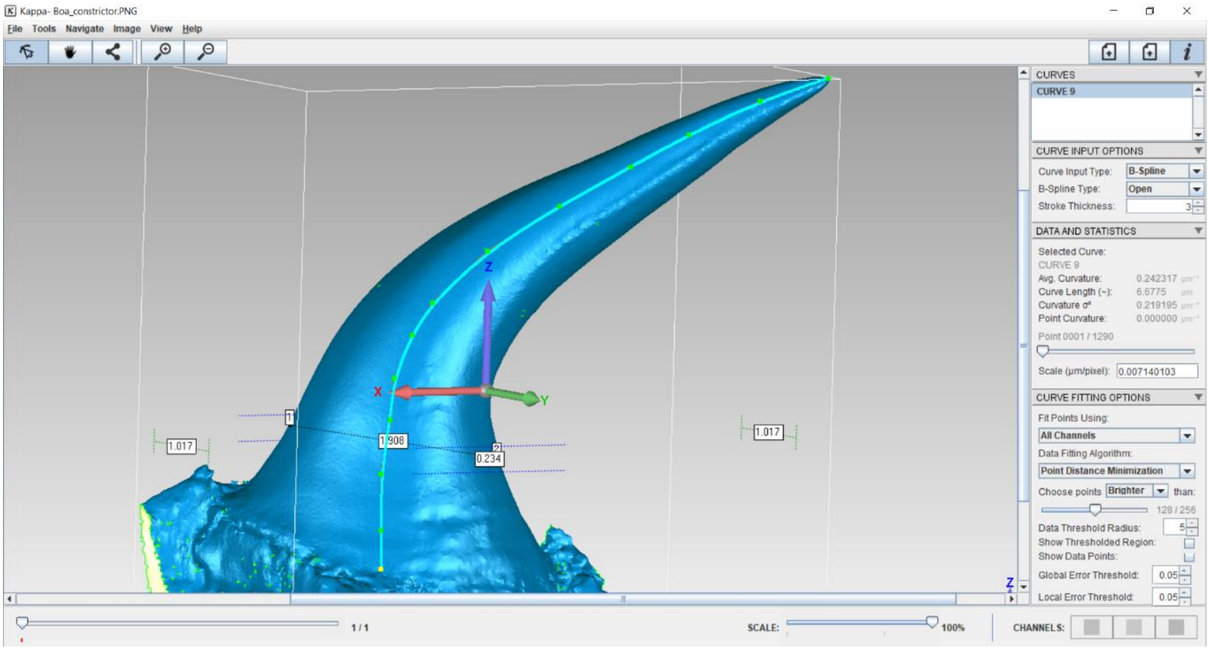


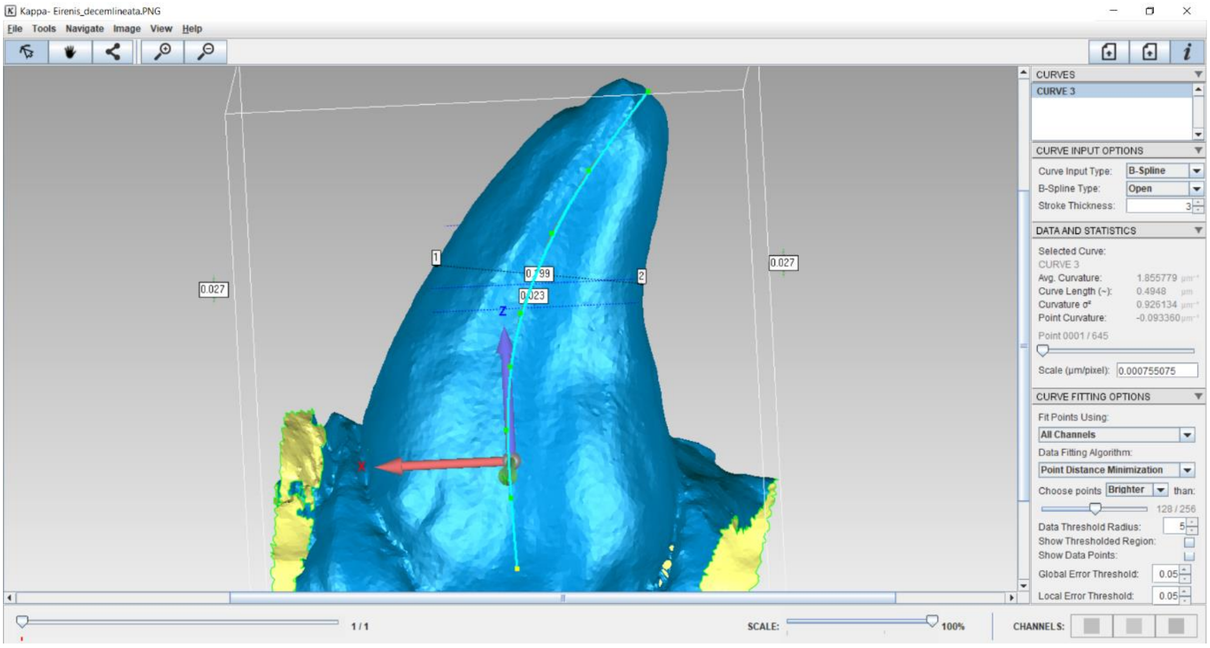
